# Expanding the Characterization of Microbial Ecosystems using DIA-PASEF Metaproteomics

**DOI:** 10.1101/2023.03.16.532922

**Authors:** David Gómez-Varela, Feng Xian, Sabrina Grundtner, Julia Regina Sondermann, Giacomo Carta, Manuela Schmidt

**Affiliations:** Systems Biology of Pain, Division of Pharmacology & Toxicology, Department of Pharmaceutical Sciences, Faculty of Life Sciences, University of Vienna, Vienna, Austria

## Abstract

Metaproteomics is gaining momentum in microbiome research due to the multi-dimensional information it provides. However, current approaches have reached their detection limits. We present a highly-sensitive metaproteomic workflow using the extra information captured by Parallel Accumulation-Serial Fragmentation (PASEF) technology. The comparison of different acquisition modes and data analysis software packages showed that DIA-PASEF and DIA-NN doubled protein identifications of mouse gut microbiota and, importantly, also of the host proteome. DIA-PASEF significantly improved peptide detection reproducibility and quantification accuracy, which resulted in more than twofold identified taxa, reaching depths comparable to metagenomic studies. Consequently, DIA-PASEF exhibited improved coverage of functional networks revealing 131 additional pathways compared to DDA-PASEF. We applied our optimized workflow to a pre-clinical mouse model of chronic pain, in which we deciphered novel host-microbiome interactions. In summary, we present here a metaproteomic approach that paves the way for increasing the functional characterization of microbiome ecosystems and is applicable to diverse fields of biological research.

## Introduction

The characterization of microbial communities (microbiomes) has increasingly gained attention due to their crucial role in the health of all planetary ecosystems^1,2^. However, our understanding of dynamic and multidimensional interactions between members of these ecosystems (host, bacteria, pathogens), and of the resulting functional alterations is still limited. Genomic sequencing methods are the most common approaches to studying these interactions by means of characterizing the taxonomic composition and relative abundance of functional genes. However, the presence of a protein-coding gene does not always result in its expression under any given condition and even transcript levels can only partially predict protein levels^3^.

Thus, prediction algorithms are used for inferring the functional roles of microorganisms in an ecosystem based on genomic or transcriptomic datasets, yet such algorithms exhibit limited performance^4,5^. Moreover, the inference on how changes in the abundance of specific taxa affect the functional output of a microbial habitat is unclear given prominent functional redundancy^6,7^. To overcome these boundaries, mass spectrometry-based metaproteomics has emerged as an attractive alternative. Metaproteomics has the potential to understand complex host-microbiome interactions as it enables the analysis of entire sets of proteins, providing direct insights into the identity and functionality of microorganisms present in an ecosystem^8^. However, current metaproteomics acquisition methods have reached their theoretical profiling limits^9^. Further advances are necessary to increase the depth and resolution of metaproteomics to facilitate a thorough investigation of microbiological habitats.

The predominant method employed for the acquisition of metaproteomic data is data-dependent acquisition (DDA) mass spectrometry. DDA generally fragments only the most intense peptide ions, rendering the majority of remaining peptides unidentifiable^10^. Additionally, accurate quantification is challenging, owing to the inconsistent recording of ion intensities along the chromatographic profile.

Consequently, reproducibility across repeated analyses is hampered and a sensitivity bias toward high-abundance peptides is introduced. These limitations are exacerbated in highly complex microbial samples (e.g., feces), which are estimated to be composed of 100 million peptide species^11^ distributed across a very high dynamic range^9^. To overcome these constraints, two recent studies have evaluated the performance of data-independent acquisition (DIA) mass spectrometry as a potential alternative method^12,13^. Although DIA offers reproducibility improvements over DDA methods, spectral complexity and sampling efficiency remain a challenge^14^, particularly in highly complex peptidomes. Consequently, the real potential of DIA-based methods in metaproteomics remains unexplored.

In our previous work^15^, we showed how the combination of the Parallel Accumulation-Serial Fragmentation (PASEF) technology^16^ and data analysis based on deep neuronal networks, incorporated in the DIA-NN software^17^, significantly increases the number of proteins quantified in mouse tissues. PASEF incorporates an additional ion mobility separation that allows for the differentiation of peptide signals that would otherwise be co-fragmented. This results in a more than tenfold increase in MS/MS scan rates without loss of sensitivity. Here, we aimed at investigating whether similar improvements can be reached in complex microbial samples, i.e., in mouse feces.

We could show that a DDA-PASEF-based workflow increased peptide identification 3-5 times and reached better data consistency and quantification reproducibility compared to previously published DDA^18^ and DIA^12^ methods. Moreover, we demonstrated that a DIA-PASEF strategy sets new profiling standards in metaproteomics by enabling the quantification of more proteins as DDA-PASEF while analyzing ten times fewer peptides. Remarkably, DIA-PASEF offered comparable taxonomical depth to previous metagenomic studies, and, at the same time, achieved a twofold increase of functional annotations compared to DDA-PASEF. When applied to a pre-clinical mouse model of neuropathic pain, DIA-PASEF enabled the quantification of more than 15,000 protein groups and deciphered novel host-microbiome interactions specific to the establishment of chronic pain. Taken together, we expect DIA-PASEF to shed light on previously unexplored regions of the metaproteome and significantly enhance our understanding of microbiological ecosystems.

## Results

### Development of a PASEF-based metaproteomic workflow

Current metaproteomic methods have reached their identification and quantification limits in complex microbial samples with high dynamic range^9^ (e.g., feces). We speculated that the extra layers of information accessible by the PASEF technology^16,19^ could offer significant improvements similar to what we demonstrated in our recent work on complex mouse tissue samples^15^. PASEF creates an extra ion mobility dimension that increases the sequencing speed ten-fold and reduces spectral complexity. As a first step of our workflow (Figure 1), we optimized the protocol for the preparation of mouse fecal samples requiring only a few manual liquid handling steps. Samples were analyzed on a timsTOF Pro mass spectrometer (Bruker Daltonics) equipped with a dual TIMS cartridge and using the PASEF acquisition method^20^. In addition to implementing PASEF using a DDA acquisition scheme (DDA-PASEF), we also tested the potential advantages of the DIA-PASEF acquisition mode^14^, which produces nearly complete data sets with peptide features defined in a four-dimensional data space (retention time, m/z, ion mobility, and intensity). Finally, we benchmarked two of the most commonly used software solutions for analyzing multi-dimensional data sets generated in DDA- and DIA-PASEF modes, MaxQuant^21^ and DIA-NN^22^.

**Figure 1.**
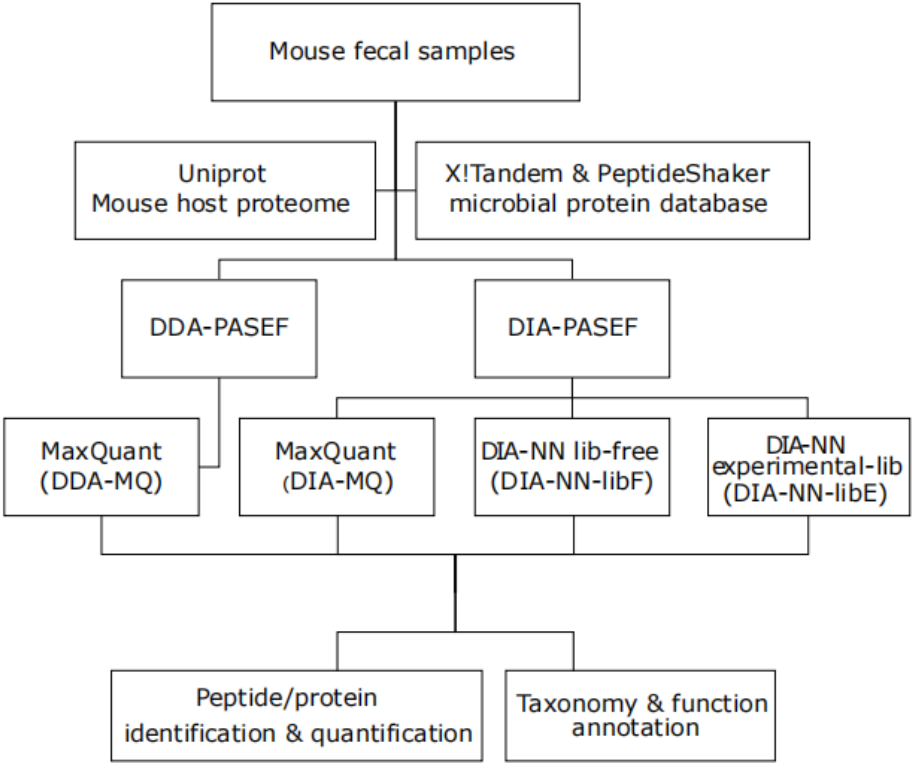
Study workflow depicting different mass-spectrometry acquisition modes and data processing steps.

### Identification and quantification performance

To compare the performance of different PASEF acquisition modes and data analysis software solutions, we first evaluated the total amount of microbial and mouse peptides and proteins, which were quantified in 10 technical replicates of a pooled mouse fecal sample (500 ng of total peptides per run). The data generated in DDA-PASEF mode were analyzed within MaxQuant (DDA-MQ), while DIA-PASEF data were analyzed using the TIMS MaxDIA module of MaxQuant (DIA-MQ) and DIA-NN. For the latter, we compared the performance employing an experimentally generated spectral library (DIA-NN-libE) to the library-free mode (DIA-NN-libF). In all cases, we set a protein and precursor FDR cutoff of 1%, as calculated by each software.

We found that DIA-NN-libE yielded nearly two-fold improvements in the number of identified microbial peptides and proteins compared to DDA-MQ. In addition, the amount of identified host peptides and proteins were also increased two-fold (Figure 2A, Supplementary Table 1). We found similar results when the library-free mode (DIA-NN-libF) was used. This performance boost resulted in a fifteen-fold increase in sensitivity as highlighted by a higher detection of microbial and host peptides and proteins by DIA-NN-libE using 31.25 ng of peptides compared to 500 ng analyzed by DDA-MQ (Figure 2B, Supplementary Table 2). Importantly, all tested DIA approaches generated data matrices with considerably fewer missing identifications for both the microbial and host proteins, compared to DDA (Figure 2C). A comparison with previous studies^12^ showed that the peptide overlap enabled by the PASEF principle in DDA mode vastly improved compared to classical DDA approaches and reached similar performance as previously published DIA strategies^12^. Remarkably, DIA-NN-libE set new standards of reproducibility compared to other methods (Figure 2D). These differences in identification performance are not due to differences in the FDR algorithms used by each software, as shown by a two-species strategy (Supplementary Figure 1). On the contrary, DIA-NN-libF detected significantly fewer false positive precursors than DDA-MQ at any given FDR level (Supplementary Figure 1).

**Figure 2.**
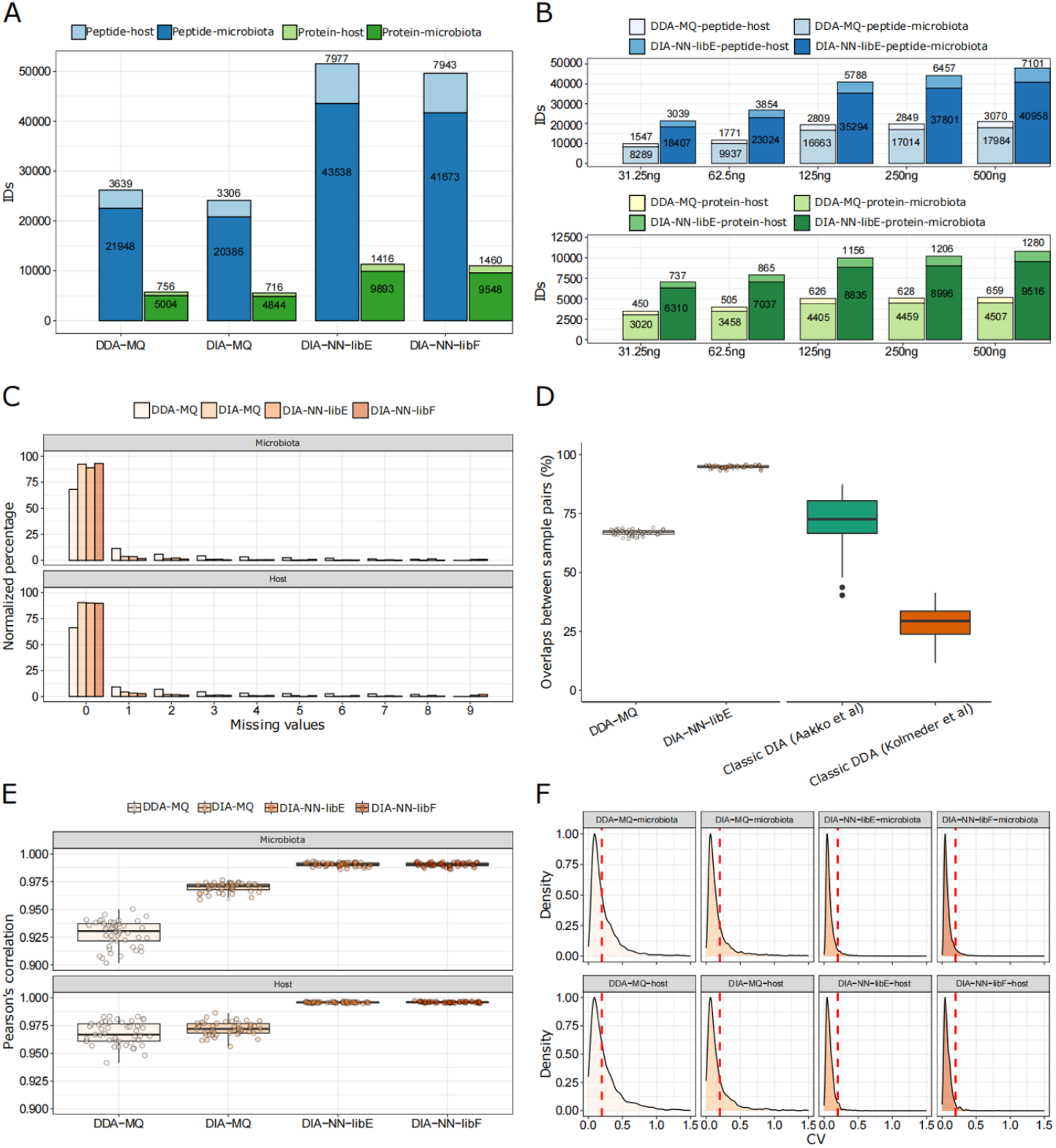
Performance evaluation of DDA- and DIA-PASEF analyzed with indicated software solutions. A) Total amount of microbial and host peptides (dark and light blue, respectively) and proteins (dark and light green, respectively) identified in 10 technical replicates of a pooled mouse fecal sample in four different workflows. B) Number of peptide (blue colors) and protein (green colors) identifications in either DDA-MQ or DIA-NN-libE workflows at different peptide injection quantities analyzed in triplicates. C) Data completeness of microbial (upper) and host (lower) protein identifications in different workflows across 10 technical replicates. D) Overlap of peptide identifications across all paired comparisons from 10 technical replicates in DDA-MQ and DIA-NN-libE workflows, and comparison with 2 previous studies. E) Intra-group correlations of 4 workflows across all paired comparisons from 10 technical replicates. The dots indicate the Pearson’s correlation for each possible paired comparison of proteins. F) Coefficient of variation (CV) distribution of proteins in the microbiota (upper) and host (lower) across 10 technical replicates in 4 workflows. The red dotted line indicates the CV = 0.2.

We further critically evaluated the quantification performance of the DDA- and DIA-PASEF approaches. DIA-NN-libE quantified 12,700-28,487 peptides, which were not detected by DDA-MQ irrespective of injected amounts (Supplementary Figure 2A). A detailed analysis showed that DIA-PASEF detected peptides covering over 4 orders of abundance magnitude, showing the biggest gain, compared to DDA-PASEF, in the lower intensity ranges (Supplementary Figure 2B). These improvements were accompanied by better protein intensity correlations among all ten technical replicates in all three DIA-PASEF-based analyses (Figure 2E), which was preserved even at the lowest sample amount tested (Supplementary Figure 3). As a result of this superior quantification accuracy, the number of proteins with a coefficient of variability (CV) < 20% significantly increased in DIA-PASEF (61% in DDA-MQ, 75% in DIA-MQ, and 94.49% in DIA-NN-libE; Figure 2F). In summary, our data clearly show the benefits of the fourth data dimension unlocked by PASEF and positions DIA-PASEF as a new state-of-the-art approach to shed light on unexplored areas of the microbial metaproteome.

### Taxonomical and functional profiling

We next assessed the performance of DDA-PASEF and DIA-PASEF in identifying microbial taxa. Globally, DIA-PASEF identified organisms belonging to the four major kingdoms present in the mouse gut microbiome (Figure 3A). Using the stringent cut-off of at least three taxon-specific peptides quantified across all analysed samples, DIA-PASEF increased the identification performance from 1.48-folds (at the species level) to 2.26-folds (at the family level) compared to DDA-PASEF (Figure 3B, Supplementary Table 3). A detailed view revealed remarkable identification gains for several phyla such as *Actinobacteria, Bacteroidetes*, and *Firmicutes* in addition to fungi and metazoans (Figure 3B). We then benchmarked our data against published metagenomic studies^23^ revealing that our workflow detected comparable numbers of mouse bacterial taxa. In general, DIA-PASEF detected genera known to constitute the core gut microbiome of healthy mice reaching a lower limit of 0.04% relative abundance, as calculated by 16S RNA sequencing^24^ (genus: *Gordonia*; Supplementary Table 4).

**Figure 3.**
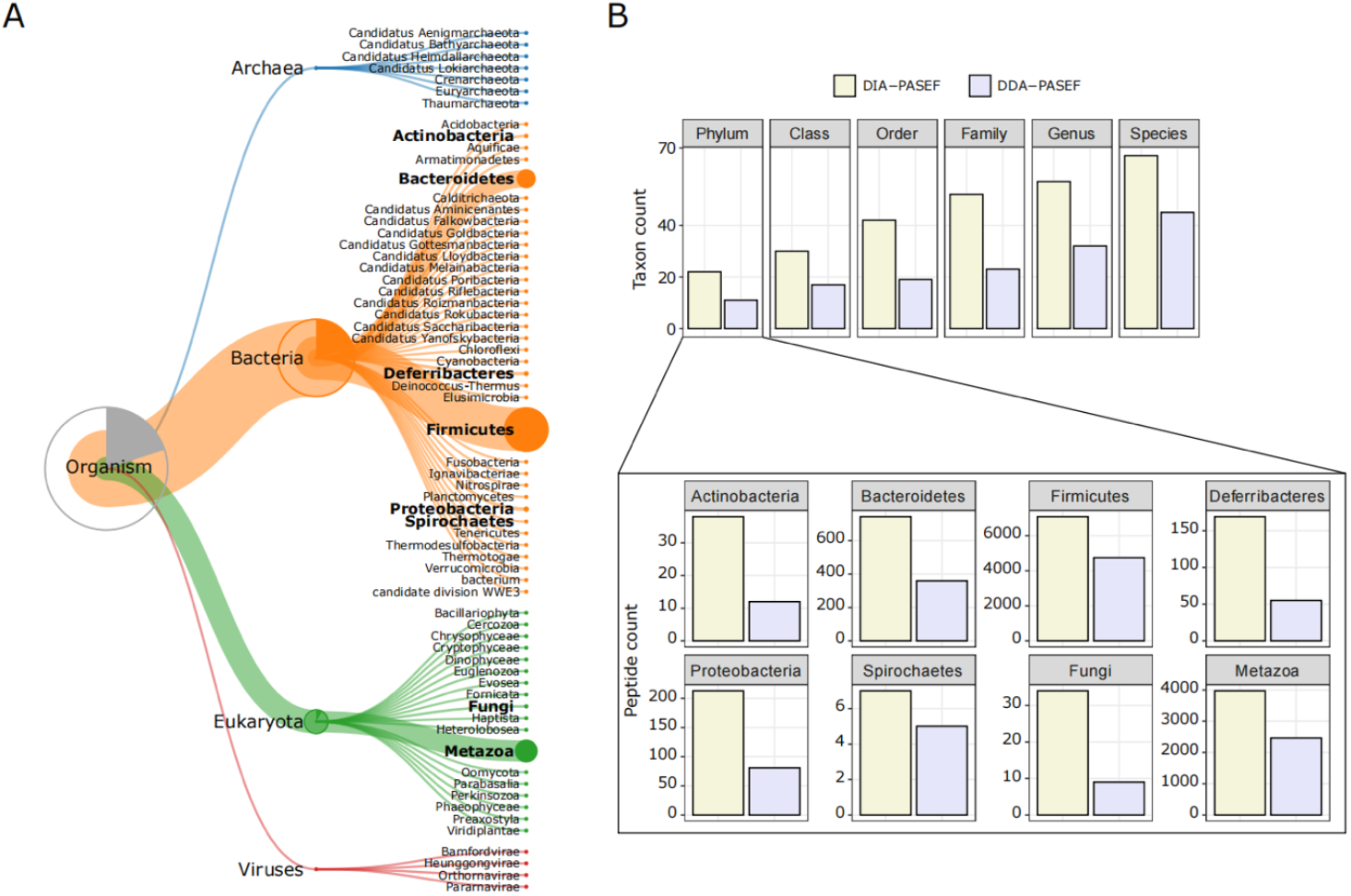
Taxonomic profiling using identified peptides from 3 technical replicates (250 ng peptide per MS run) in DDA-MQ or DIA-NN-libE workflows. A) Hierarchical classification of annotated taxonomy from Unipept using DIA-NN-libE data. B) Comparison of DDA-PASEF (lavender) and DIA-PASEF (beige) workflow at different taxonomic levels (upper panel: cut-off: at least 3 unique peptides per taxon as identified in iMetalab). The zoom-in bar plots show phyla annotated in both workflows with their respective peptide counts.

Next, we investigated the annotation of detected proteins to functional pathways for both the microbiota and the host. The aforementioned improvements in protein quantification reached by DIA-PASEF translated to increased numbers of KEGG, COG, and NOG functional terms compared to DDA-PASEF (Figure 4A). DIA-PASEF data covered 100% of all KEGG pathways detected by DDA-PASEF and enabled the quantification of additional 54 pathways (Supplementary Table 5). We also compared pathways defined by exclusive proteins (i.e., proteins that are not shared among different KEGG terms) and found that DIA-PASEF offered higher protein coverage in almost all common pathways and identified unique ones (Figure 4A).

**Figure 4.**
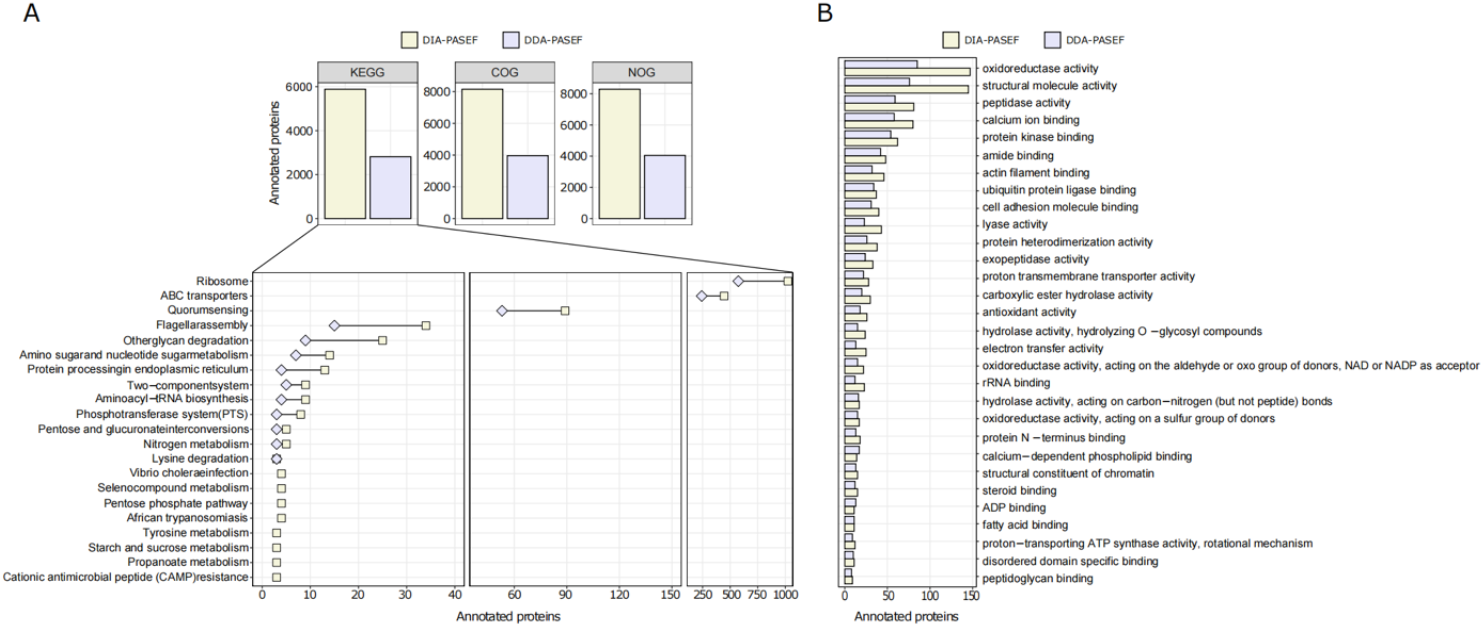
Functional annotations of metaproteomes using DDA-PASEF (lavender) and DIA-PASEF (beige) workflows in 3 technical replicates (250 ng peptide per MS run). A) Bar plots show the number of microbial proteins annotated using KEGG, COG, and NOG databases. The zoom-in lower panel depicts KEGG pathways with uniquely annotated microbial proteins (cutoff: at least 3 proteins per pathway). B) Top 30 common GO-MF (Gene Ontology-Molecular Function) enriched in the host proteome (analyzed by Metascape using a cut-off of 3 proteins per function and an adjusted p value < 0.01).

Noteworthy, the latter subset is mainly related to the metabolic processing of compounds such as tyrosine, sucrose, and propanoate (Figure 4A). As for the host biology, DIA-PASEF enabled the characterization of 77 unique protein functional categories (Supplementary Table 6), as well as an increased number of host proteins associated with Gene Ontology Molecular function terms (GO-MF) commonly detected by DDA-PASEF (Figure 4B).

### DIA-PASEF offers novel insights into host-microbiome interactions upon neuropathic pain

Chronic pain affects an estimated 20% of the world’s population^25^. Nevertheless, current therapies have severe limitations and significant side effects^26^, such as addiction and overdose. Initial studies utilizing 16S RNA sequencing showed changes in the gut microbiome composition during the onset and regulation of chronic pain in humans and animal models^27^, suggesting the gut microbiome as a potential target for novel treatments. However, our knowledge is scarce and a thorough assessment of distinct functional alterations of the gut host-microbiome relevant for chronic pain is awaiting.

We set out to test our metaproteomic workflow in the spared nerve injury (SNI) mouse model of chronic neuropathic pain^28^. We collected fecal samples of aged-matched SNI, SHAM, and Naive female mice before (Pre) and 14 days (14D) post-surgery covering the critical period of chronic pain onset^29,30^ (Figure 5A). Analysis of mouse behaviours confirmed the expected development of neuropathic pain as indicated by mechanical allodynia^30^ (i.e., hypersensitivity to an innocuous tactile stimulus to the affected hind paw) in SNI but not in SHAM mice (Supplementary Figure 4 and Supplementary Table 7). In total, our DIA-PASEF workflow detected 81,378 peptides corresponding to 12,239 and 2,267 microbial and mouse protein groups, respectively (Figure 5A, Supplementary Table 8). We quantified 29 phyla, 45 classes, and 119 species with at least 3 peptides per taxon (Supplementary Figure 5, Supplementary Table 9). In comparison to previous metagenomic studies^31,32^, we quantified more and unique low-abundant taxa. Overall, we observed a similar abundance distribution as a recent study using 16S RNA sequencing in fecal samples of 600 mice^31^ (Supplementary Table 10). Moreover, the sensitivity of DIA-PASEF enabled us to detect species with a mean abundance of 0.003 (when using the cutoff of 3 species-specific peptides, e.g., *Alistipes indistinctus*) or 0.00007 (when using the cutoff of 1 species-specific peptide, e.g. *Paenibacillus amylolyticus*) as calculated from a recent compilation of 2,446 global mouse gut metagenomes^32^ (Supplementary Table 10).

**Figure 5.**
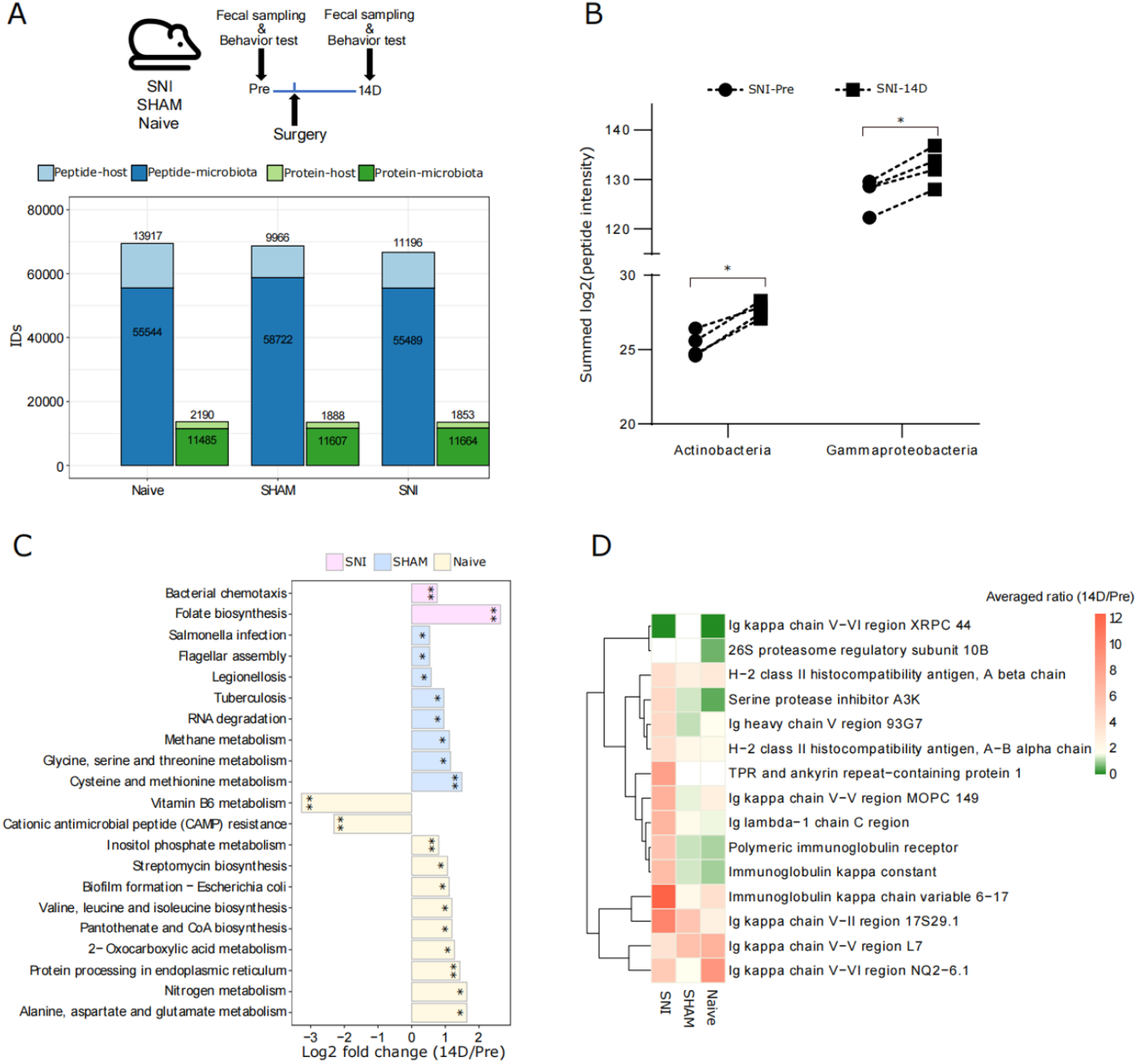
Host-microbiome changes in the mouse gut during neuropathic pain. A) *Top*: Experimental workflow indicating the three experimental groups and the times of fecal sampling, surgery procedures, and behavioral assessment (for details, please see Methods). *Bottom*: Peptide and protein identifications in both the microbiota and the host, using DIA-PASEF in all three experimental groups (Naive, SHAM, and SNI). B) The abundance of the two indicated bacterial classes that are specifically up-regulated in SNI mice at 14D post-surgery (q value <0.05; paired t-test with Benjamini-Hochberg correction for multiple comparisons). C) Significantly altered KEGG pathways of the microbiome in all three experimental groups (* q<0.05, ** q<0.01). D) Expression patterns of 15 host proteins (selected based on p<0.005, pair-wise comparison 14D versus Pre) in all three experimental

Interestingly, statistical analysis revealed that the bacterial classes of *Actinobacteria* and *Gammaproteobacteria* are exclusively increased during neuropathic pain in SNI animals at 14D (Figure 5B; Supplementary Table 11).

To gain insights into host-microbiome interactions and their functional changes upon nerve injury, we investigated proteins and their association with biological pathways. Pathway analysis using KEGG terms showed significant enrichment of distinct microbial pathways in each of the three experimental conditions when comparing 14D with “Pre” data (Figure 5C; Supplementary Table 12). At the host level, all three conditions showed only limited overlap but rather quite specific pathway alterations (Supplementary Figure 6A): while in SNI animals there was a clear upregulation of processes related to the immune response of the gut (Figure 5D), in SHAM animals the repertoire of altered proteins was distinct, and the extent of observed changes was significantly lower except for the increase of several major urinary proteins (Supplementary Figure 6B). Our results provide a stepping stone for follow-up interventional studies investigating the mechanistic relevance of here discovered host-microbiome alterations for neuropathic pain and its chronification. Besides these novel findings in experimental groups (SNI and SHAM), our data reveal - for the first time – prominent functional changes occurring in the gut of age-matched Naive mice between 4 and 6 weeks of age i.e., a period of 14 days during which our experiments were conducted (Figure 5A). Thus, our results highlight the need to adequately control for purely age-induced changes in microbiome studies by including Naive non-treated mice.

## Discussion

We present a novel metaproteomic workflow utilizing PASEF technology, as well as the deep neural network-based data analysis offered by DIA-NN. The combination of both elements rendered very significant improvements compared to previous metaproteomic workflows and facilitated, for the first time, the global assessment of host-microbiome interactions in a mouse model of neuropathic pain.

We hypothesized that the features of the PASEF technology would be especially advantageous in a sample with high dimensionality like the fecal microbiome, where few species represent more than 80% of the total abundance^9^. The sequential release of ion packages trapped in the tunnel of the TIMS mobility analyzer together with the synchronization of the quadruple selection increases sensitivity and reduces the complexity of spectra generated by PASEF^16^. Accordingly, we show that the reproducibility of detection between samples offered by DDA-PASEF doubled compared to classical DDA approaches and even matched the performance of classical DIA^12^. Adding a DIA mode to the PASEF scheme enabled us to significantly increase the number of peptides and proteins quantified in comparison to DDA-PASEF. These gains became overtly evident in peptides at the lower abundance range, a fact, that will greatly improve the detection limits of current metaproteomic workflows^9^. In this line, DIA-PASEF reached similar or even better taxonomical depth as previous large-scale mouse metagenomic studies. Upcoming DIA-PASEF schemes^33-35^ have the potential to further advance the impact of PASEF on metaproteomics that we pioneered in this study.

DIA-PASEF achieves a nearly complete record of all peptide fragments^14^. Despite the significant improvements shown in our study, a large amount of peptide spectra likely remains unidentified. However, this information is stored in very comprehensive digital MS/MS datasets. Thus, our workflow offers the possibility to interrogate these by either using continuously improved microbiome-based databases or emerging AI-based algorithms. The use of these datasets in the metaproteomic field will offer extra advantages: community data resources, optimization in the utilization of valuable microbiome samples (e.g., clinical human samples), and reduced animal experimentation for hypothesis testing.

The taxonomical and functional depths achieved by DIA-PASEF allowed us to directly interrogate, for the first time, the functional alterations in the gut ecosystem during neuropathic pain. Our findings suggest that *Actinobacteria* and *Gammaproteobacteria* are exclusively increased upon neuropathic pain in SNI animals. These data are in line with previous genomic findings in SNI mice^36^ and interstitial cystitis human pain syndrome^37^.

Interestingly, we detected an upregulation of the microbial KEGG pathway “folate biosynthesis” exclusively upon nerve injury. Folate is produced in the murine gut by several bacterial taxa including *Actinobacteria*^38^, and its deficiency has been associated with a greater risk of peripheral neuropathy in a retrospective cohort of more than 500,000 people^39^. Remarkably, Botez and colleagues^40^ showed an improvement in clinical and electrophysiological measurements of five patients with polyneuropathy after 9-39 months of folate therapy. Therefore, the functional insights offered here might indicate a mechanism by which the gut ecosystem aims at ameliorating neuropathic pain. This is an exciting hypothesis that may open new therapeutical avenues harnessing folate supplementation as implemented for diverse autoimmune diseases^41^.

Moreover, folate stabilises and increases the abundance of Treg lymphocytes in the colon^41^. Interestingly, T_reg_ lymphocytes alleviate chronic neuropathic pain by inhibiting the Th1 response via CD4+ helper cells^42^. Emerging clinical evidence proposes that the pathogenesis of neuropathic pain may be a centrally-mediated neuroimmune phenomenon with an important contribution by the gut microbiome ^43^, e.g. by influencing the balance between pro-inflammatory and anti-inflammatory T cells^44^.

In addition to the potential role of bacterial folate production, we also discovered alterations of several host protein complexes, exclusively in SNI animals. For example, those known to be involved in the crosstalk between the host immune system and the gut microbiome. We observed an upregulation of two components of the Major Histocompatibility Complex II (MHC-II). MHC-II contributes to the maturation of B cells via the presentation of exogenous bacterial antigens and the production of secreted IgA^45^(SIgA). Dysregulation of the IgA-microbiota axis affects multiple pathologies with an inflammatory component^46^ but its link to pain syndromes is unknown. Notably, we observed a significant upregulation of the Polymeric Immunoglobulin Receptor in SNI animals at 14D. Polymeric Immunoglobulin Receptor represents one of the two proteins that are necessary for the secretion of IgA from gut mucosal plasma cells into the gut lumen where it contributes to controlling the abundance of commensal microbiota^47^. A detailed analysis of our data shows that both the immunoglobulin J chain (J chain), also being part of this secretory complex, as well as IgA tend to be upregulated in SNI animals, even though they did not pass our statistical cutoff (P01592 and P01878 in Supplementary Table 13). In addition, we observed a significant increase in *Gammaproteobacteria* in SNI mice. In this context, it is noteworthy that *Gammaproteobacteria* are known to produce gut inflammation in mice thereby triggering an IgA-dependent immune response via activation of B cells^48^. Thus, our findings open new avenues for future mechanistic studies aimed at deciphering the role of these microbiome-immune pathways for chronic pain.

In summary, we present the enormous potential of DIA-PASEF to provide in-depth and unprecedented insights into host-microbiome interactions with a significant impact on our understanding of microbiological ecosystems across diverse biology disciplines.

## Method and Material

### Reagents

All reagents were purchased from Sigma-Aldrich (St. Louis, Missouri) if not mentioned otherwise. Acetonitrile (ACN) and formic acid (FA) were purchased from Fisher Scientific (Hampton, New Hampshire; both FA and ACN were liquid chromatography-mass spectrometry (LC-MS) grade). LC-MS grade water from Sigma was used for all solutions. Protease inhibitor was purchased from Roche (Complete Ultra Tablets Mini, Roche, Basel, Switzerland).

### Animal housing and surgery

In-house bred C57BL/6J female mice were used. All animal experiments were carried out with the approval of the IACUC at the University of Vienna and of the Austrian Ministry for Education, Science and Research (BMBWF; license number 2021-0.138.925). All mice used in this study were group-housed in the same room, with a 12 h light/dark cycle, and with water and food ad libitum. All mice were weaned between 22-24 days after birth. The spared nerve injury (SNI) paradigm was used to induce neuropathic pain in female mice at 4 weeks of age. Mice were anaesthetized using isoflurane (4% for induction first and 2% for maintenance in O_2_). To expose the trifurcation of the left sciatic nerve, the skin and underlying biceps femoris muscle were incised. A ligature was placed around the common peroneal and tibial nerves with a 7-0 surgical braided silk (Vömel, Kronberg, Germany) below the bifurcation, followed by distal transection of both nerves. A 2 mm segment was excised from both nerves to impede regrowth. The skin was closed with one surgical clip (AutoClip®, Fine Science Tools, Heidelberg, Germany) and disinfected with povidone-iodine (Mundipharma, Frankfurt am Main, Germany). The operated mice were injected with carprofen (0.05mL/10g body-weight, Zoetis Österreich GmbH, Austria) immediately before surgery and on postoperative day 1. The clips were removed on postoperative day 5 with an AutoClip® remover (Mikron Precision Inc, Biel, Switzerland). In SHAM-operated mice, the same surgical procedure was carried out as in SNI but without ligating and transecting the branches of the sciatic nerve. All operated mice were weighed daily until postoperative day 7 and from then on, every second day until postoperative day 14 by female experimenters only.

### Mechanical sensitivity test

The test was performed 3-4 days before surgery (Pre) and on a postoperative day 14 (14D) using a dynamic plantar aesthesiometer (automated von Frey filament, Dynamic Plantar Aesthesiometer: 37450-001, Dynamic Plantar Aesthesiometer Touch Stimulator: 37400-002, Ugo Basile®, Gemonio, Italy). Mechanical force (Force Intensity: 10.0g, Ramp Time: 40sec) was applied to the lateral side of the plantar hind paw and the withdrawal latency (in seconds) was measured five times for each hind paw, with at least a 2-minute recovery period between each measurement. The test was done only by female experimenters.

### Fecal collection

Mice were placed into 10×10cm boxes covered with a lid, for 15min. After removing mice from the boxes, feces were collected using forceps that were previously cleaned with 70% ethanol between each sample. Samples were transferred to precooled 2mL autoclaved tubes (Eppendorf, Hamburg, Germany) and stayed on ice until stored at - 80°C.

### Protein extraction, SP3-assisted protein digestion, and peptide clean-up

For protein preparation, 30 mg of fecal samples were mixed with 200 μL of lysis buffer (5% SDS, 2M Urea, 50mM Tris-HCl, Protease Inhibitor 1x) and vortexed vigorously for 1 min. Following, samples were placed in a Thermomixer (Serial Number: 5382JR638726, Eppendorf, Hamburg, Germany) for 15 minutes at 1,200 rpm and 70°C. When finished, samples were ultrasonicated using a Bioruptor Pico (Diagenode, Seraing, Belgium; program: 15 cycles of 30 seconds “ON” and 30 seconds OFF, frequency level: Low, water temperature: +20°C). Afterwards, tubes were centrifuged at 16,000 g for 5 minutes at room temperature. Supernatants were saved in new tubes and submitted to a new round of centrifugation. The saved supernatants were stored at -80°C until SP3-assisted protein digestion.

For protein clean-up and digestion, a modified version of the single-pot, solid-phase-enhanced sample preparation (SP3) method from Hughes et al. was used^49^. Briefly, 50 μg protein was taken into a 1.5 mL LoBind tube (Eppendorf, Hamburg, Germany), and the sample volume was added up to 50 μL with lysis buffer. The fecal sample was subjected to protein reduction (5 mM Dithiothreitol, DTT, 30 minutes incubation at +60°C) and alkylation (20 mM Iodoacetamide, IAA, 30 minutes at room temperature in the dark). The remaining IAA in the sample was quenched with the addition of DTT to a final concentration of 5 mM. 10 μL of pre-mixed Sera-Mag SpeedBead beads (50 mg/ml, Cytiva, Marlborough, Massachusetts) were added to 50 μg protein sample. To initiate the binding of proteins to the beads, one volume of absolute ethanol was added immediately, followed by incubation on a Thermomixer (Eppendorf) at 24°C for 5 minutes with 1,000 rpm agitation. The supernatant was removed after 2 minutes resting on a magnetic rack, and the beads were rinsed three times with 500 μL of 80% ethanol. Rinsed beads were reconstituted in 50 μL digestion buffer (50 mM ammonium bicarbonate, pH 8). Protein digestion was performed with 2 μg of sequencing-grade trypsin/LysC (Promega, Madison, USA) for 18 hours at 37°C with 950 rpm agitation. After digestion, ACN was added to each sample to a final concentration of 95%. Mixtures were incubated for 8 minutes at room temperature and then placed on a magnetic rack for 2 minutes. The supernatant was discarded, and the beads were rinsed with 900 μL of 100% ACN. Rinsed beads were reconstituted in 20 μL LC-MS grade water to elute the peptides. The peptide concentration was measured in duplicate using NanoPhotometer N60 (Serial number: TG2022, Implen, Munich, Germany) at 205 nm. Peptide samples were acidified with FA to a final concentration of 0.1% and stored at -20°C until LC-MS/MS analysis.

### LC-MS/MS setup

Nanoflow reversed-phase liquid chromatography (Nano-RPLC) was performed on a NanoElute system (Bruker Daltonik, Bremen, Germany). Peptides were separated with either a 70 minutes or 130 minutes gradient on a 25 cm x 75 μm column packed with 1.6 μm C18 particles (IonOpticks, Fitzroy, Australia). Mobile solvent A consisted of 2% ACN, 98% water, 0.1% FA and mobile phase B of 100%, 0.1% FA. For both gradient lengths, the flow rate was set to 400 nL/min for the first 2 minutes and the last 9 minutes of the gradient, while the rest of the gradient was set to 250 nL/min. In the 70 minutes separation, the mobile phase B was linearly increased from 0 to 20% from 3 minutes to 50 minutes, flowed by a linear increase to 35% within 10 minutes and a steep increase to 85% in 0.5 minutes. Then a flow rate of 400 nL/min at 85% was maintained for 9 minutes to elute all hydrophobic peptides. In the 130 minutes separation, the mobile phase B was linearly increased from 0 to 20% from 3 minutes to 110 minutes, flowed by a linear increase to 35% within 10 minutes and a steep increase to 85% in 0.5 minutes. Then a flow rate of 400 nL/min at 85% was maintained for 9 minutes to elute all hydrophobic peptides. NanoElute LC was coupled with a hybrid TIMS quadrupole TOF mass spectrometer (timsTOF Pro, Bruker Daltonik, Bremen, Germany) via a CaptiveSpray ion source. Samples were analyzed in both data-independent acquisition (DIA) and data-dependent acquisition (DDA) modes coupled with parallel accumulation serial fragmentation (PASEF) for methods comparison. The samples used for Figure 5 (SNI, SHAM and Naive) were analyzed in DIA-PASEF. In both acquisition modes, the TIMS analyzer was operated in a 100% duty cycle with equal accumulation and ramp times of 100 ms each. Specifically, in DDA-PASEF mode^20^, 10 PASEF scans were set per acquisition cycle with ion mobility range (1/k0) from 0.6 to 1.6, and singly charged precursors were excluded. Dynamic exclusion was applied to precursors that reached a target intensity of 17500 for 0.4 minutes. Ions with m/z between 100 and 1700 were recorded in the mass spectrum. In DIA-PASEF mode^14^, precursors with m/z between 400 and 1250 were defined in 16 scans containing 32 ion mobility steps with an isolation window of 26 Th in each step with 1 Da overlapping for neighbouring windows. The acquisition time of each DIA-PASEF scan was set to 100 ms, which led to a total cycle time of around 1.8 sec. In both DDA- and DIA-PASEF modes, the collision energy was ramped linearly from 59 eV at 1/k0 = 1.6 to 20 eV at 1/k0 = 0.6.

### Protein database generation for metaproteome analysis

A metagenome-translated protein database (PD1) was downloaded from http://gigadb.org/ containing 2.6 million protein sequences^50^. Due to the large size of the protein database, the following approach was applied to generate a reduced and sample-specific protein database. Briefly, a fecal pooled peptide sample was recorded 10 times using DDA-PASEF and a 70 minutes chromatography gradient. The 10 raw data files were first converted into .mgf format and then searched against the PD1 using X!Tandem^51^, a search engine integrated into SearchGUI^52^ (Version 4.1.11). Trypsin was specified with a maximum of 2 missed cleavages allowed. The search included variable modifications of methionine oxidation and N-terminal acetylation and a fixed modification of carbamidomethyl on cysteine. The mass tolerances of 10 ppm for both precursor and fragment were used. The output of the X!Tandem search was further validated in PeptideShaker^53^. All validated proteins were exported as the reduced protein database (PD2) containing 9,750 protein sequences. PD2 was further used for the comparison of different workflows. Another pooled peptide sample made from Naive, SHAM and SNI mice was submitted to a 130-minute gradient and analyzed in DDA-PASEF mode in 9 replicates. The 9 raw data files were subjected to the same procedures aforementioned to generate another reduced protein database (PD3) containing 10,859 protein sequences. PD3 was further used to analyze the metaproteome of Naive, SHAM and SNI mice. Mus musculus reference proteome (PD4) was downloaded from Uniprot (https://www.uniprot.org/proteomes/UP000000589) and used to identify host proteins from the fecal samples.

### Data processing using different workflows

DDA-PASEF raw data files were analyzed with MaxQuant (version 2.1.3.0) and searched with Andromeda against PD2 and PD4. The search type was specified as “TIMS-DDA”. The minimal peptide length was set to 7 amino acids, and a maximum of 2 missed cleavages was allowed. The search included variable modifications of methionine oxidation and N-terminal acetylation, deamidation (N and Q) and fixed modification of carbamidomethyl on cysteine, and a maximum of 5 modifications per peptide was allowed. The “Match between runs” function was checked within a 0.7 min retention time window and 0.05 ion mobility window. Mass tolerance for peptide precursor and fragments were set as 10 ppm and 20 ppm, respectively. The FDR was set to 0.01 at the precursor level and protein level. Label-free quantification algorithm with a minimum 1 LFQ ratio count was used to quantify identified proteins. The rest of the parameters were kept as default.

DIA-PASEF raw data files were analyzed with MaxQuant (version 2.1.3.0) and the search type was specified as “TIMS MaxDIA”. The data were searched against the spectra library generated with the DDA-PASEF data search in the MaxQuant workflow mentioned above using the Andromeda algorithm. Specifically, the output evidence.txt, peptide.txt and msms.txt files were used. The rest of the configurations were set as same as the search for DDA-PASEF data.

DIA-PASEF raw data files were also searched in DIA-NN^17^. DIA-NN (version 1.8.1) was used to process DIA-PASEF data in library-free mode with PD2 or PD3 and PD4 to generate the predicted spectrum library. A deep learning-based method was used to predict theoretical peptide spectra along with their retention time and ion mobility. Trypsin/P was used for in silico digestion with an allowance of a maximum of 2 missed cleavages. Variable modifications on peptides were set to N-term methionine excision, methionine oxidation and N-term acetylation, while carbamidomethylation on cysteine was a fixed modification. The maximum number of variable modifications on a peptide was set to 5. Peptide length for the search ranged from 7 to 52 amino acids. Aligned with the DIA-PASEF acquisition method, m/z ranges were specified as 400 to 1250 for precursors and 100 to 1700 for fragment ions. Both MS1 and MS2 mass accuracy were set to automatic determination. Protein inference was set to “Protein names (from FASTA)” and the option of “Heuristic protein inference” was unchecked. Match-between-run (MBR) was checked for cross-run analysis. RT-dependent cross-run normalization and Robust LC (high precision) options were selected for quantification. In addition, the same DIA-PASEF raw data files were processed with DIA-NN (version 1.8.1) using an experimental spectrum library generated in our lab with several metaproteome studies. The experimental library contains microbial and host proteins of in total 14,711 protein isoforms, 19,153 protein groups and 98,159 precursors. Besides the use of the experimental spectrum library, the rest of the search parameters were the same as indicated above for the DIA-PASEF data search in library-free mode.

### Metaproteome data batch correction and normalization

For the neuropathic pain experiment, precursor intensities (Normalized Intensity from the DIA-NN main report table) were submitted to median normalization and quantile batch correction using the proBatch R package^54^. The resulting precursor intensities were further processed with the R package, DIA-NN (https://github.com/vdemichev/diann-rpackage), to extract and calculate the MaxLFQ^55^ quantitative intensity for all identified peptides and protein groups with q-value < 0.01 as criteria at precursor and protein group levels.

### Taxonomy and function annotation

iMetaLab^56^ (version 2.3.0) was used for taxonomy and function annotation. The MaxLFQ peptide data (microbial and host) were imported into iMetaLab, and the built-in taxonomy database was used for the mapping (Ignore blanks below rank: Superkingdom, Unique peptide count >=3). MaxLFQ protein identifications with corresponding intensities were imported into iMetaLab for functional annotation (using default parameters).

### Taxonomic quantification strategy

The iMetalab’s taxonomy annotation output underwent further processing in R, where the following steps were taken to quantify the detected taxa in each sample. Firstly, peptides without annotations at a specific taxon level (e.g., Species) were removed. Next, taxa with at least three unique peptides annotated were retained. The peptide intensity in each sample was then extracted based on the retained taxa. Peptides that were not quantified in all samples were removed. Finally, the log2-transformed intensities of common peptides annotated in the same taxon were summed up in each sample. The intensities matrices of 14D and Pre in each condition were subjected to paired t-test in GraphPad Prism (version: 9.5.0(730)) using the “Benjamini and Hochberg” method for multiple comparisons.

### Functional quantification strategy

The output of the function annotation in iMetaLab underwent further processing in R using the following steps to calculate the intensity of each KEGG pathway in each sample using proteins with a significance p < 0.005 when comparing 14D versus Pre within each condition (paired t-test in Limma package^57^). Firstly, pathways annotated with the same protein were separated into rows with duplicated intensities for this protein in all samples, as there is no evidence to suggest that the protein belongs to only one pathway. Secondly, the intensities of proteins annotated in the same pathway were summed up in each sample. Thirdly, the number of proteins mapped to each pathway in each sample was counted. Finally, the intensity of each pathway was normalized by dividing the summed intensity by the number of proteins mapped. The intensity of each pathway was log2-transformed before statistical analysis. The transformed intensities were tested by comparing 14D versus Pre in each condition (Naive, SHAM, and SNI) using the Limma^57^ R package in a paired manner with the “Benjamini and Hochberg” method for multiple comparisons.

### Pathway analysis of host proteome

The host protein identifications were analyzed in Metascape^58^ (https://metascape.org/, version 3.5.20230101, accessed on 2023.02.17) and searched against the Gene Ontology Molecular Function database with default settings.

## Acknowledgement

This work is funded by a Research grant from the FWF (P35856-B to MS) and the University of Vienna. We would like to thank Barbara Berger and Nicole Schranzer (Division of Pharmacology & Toxicology, University of Vienna, Austria) for their excellent technical assistance. We are grateful to Hanna Fischer (Division of Pharmacology & Toxicology, University of Vienna, Austria) for short-term help with mouse behaviour, to Nicole Kanta for help with video-analysis, to the Systems Biology of Pain team at the University of Vienna for discussions, to Daniel Segelcke (University Hospital Muenster, Muenster, Germany) for advice on mouse surgeries, and to Kai Cheng for his assistance with iMetalab.

## Data availability

The DDA- and DIA-PASEF raw files, and the experimental spectral library have been deposited to the PRIED database (https://www.ebi.ac.uk/pride/) and can be found by searching for the title of the manuscript.

## Ethics

All animal experiments were carried out in strict accordance with institutional IACUC guidelines, international ARRIVE guidelines, and the principles of the 3Rs of animal research in accordance with the BMBWF (Federal Ministry of Education, Science and Research, Austria). All experimental procedures were approved by the BMBWF (GZ 2020-0.592.919).

## Author contributions

Conceptualization: D.G.V.; experimental design: D.G.V.; biochemistry and mass spectrometry: F.X. and D.G.V.; data analysis: F.X., D.G.V., S.G., and G.C.; animal experiments: S.G. and J.S.; writing initial draft: D.G.V.; writing: all authors edited and approved the final manuscript; supervision: D.G.V. and M.S.; Project administration: DGV, MS; Funding acquisition: MS.

## Figures

**Supplementary Figure 1:**
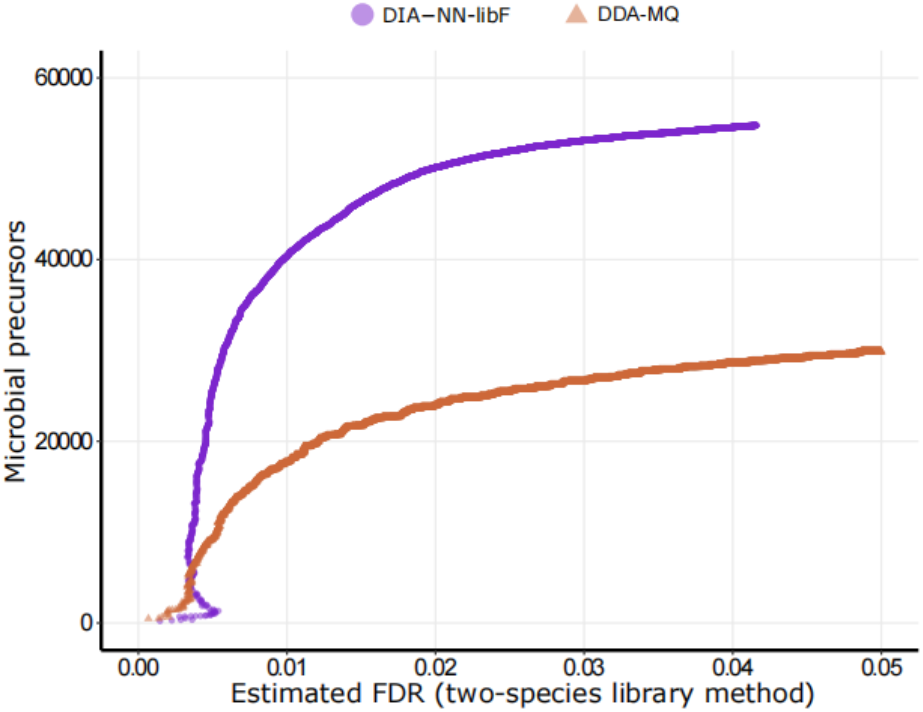
Identification performance of DIA-NN and MaxQuant software. Precursor identification numbers are plotted against the FDR, estimated using a two-species library method, searching the data against both the microbial protein database (PD2) and the A. Thaliana proteome (UP000006548). Each point in the graph corresponds to a decoy (*A*.*Thaliana*) precursor with the x-axis reflecting its estimated FDR and the respective score threshold and the y-axis representing the number of target microbial precursors at this threshold.

**Supplementary Figure 2:**
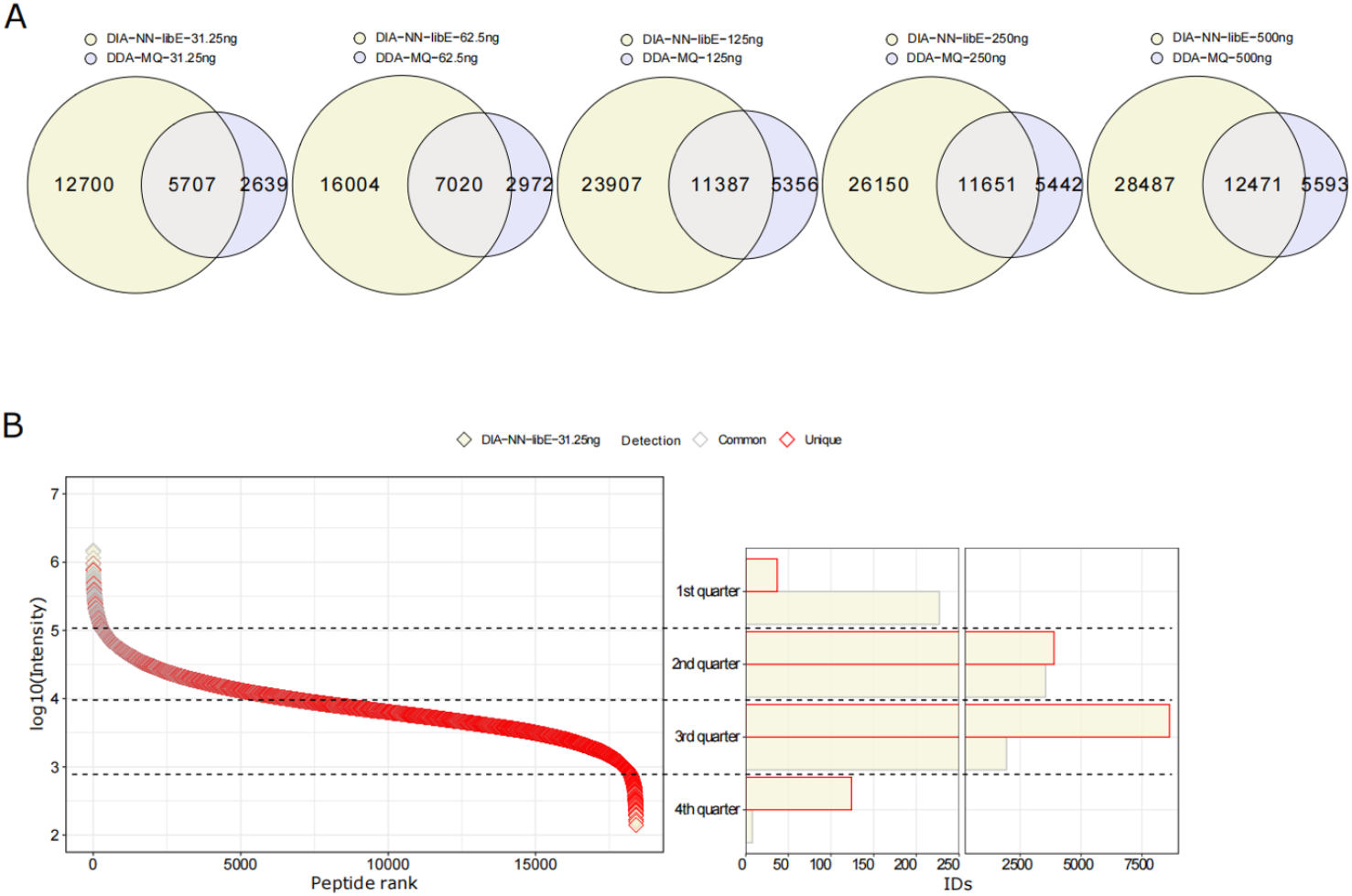
Comparisons of peptide identifications. A) Venn diagrams show the unique and overlapped number of identified microbial peptides between the DIA-NN-libE and the DDA-MQ workflows, at increasing amounts of the injected peptides. B) *Left*: Dynamic range of peptides uniquely identified using DIA-NN-libE (red symbols) or shared with DDA-MQ (grey symbols), when 31.25 ng of peptides were injected. *Right*: Number of detected peptides in each intensity quarter either exclusively identified using DIA-NN-libE (red-bordered boxes) or shared with DDA-MQ (grey-bordered boxes).

**Supplementary Figure 3:**
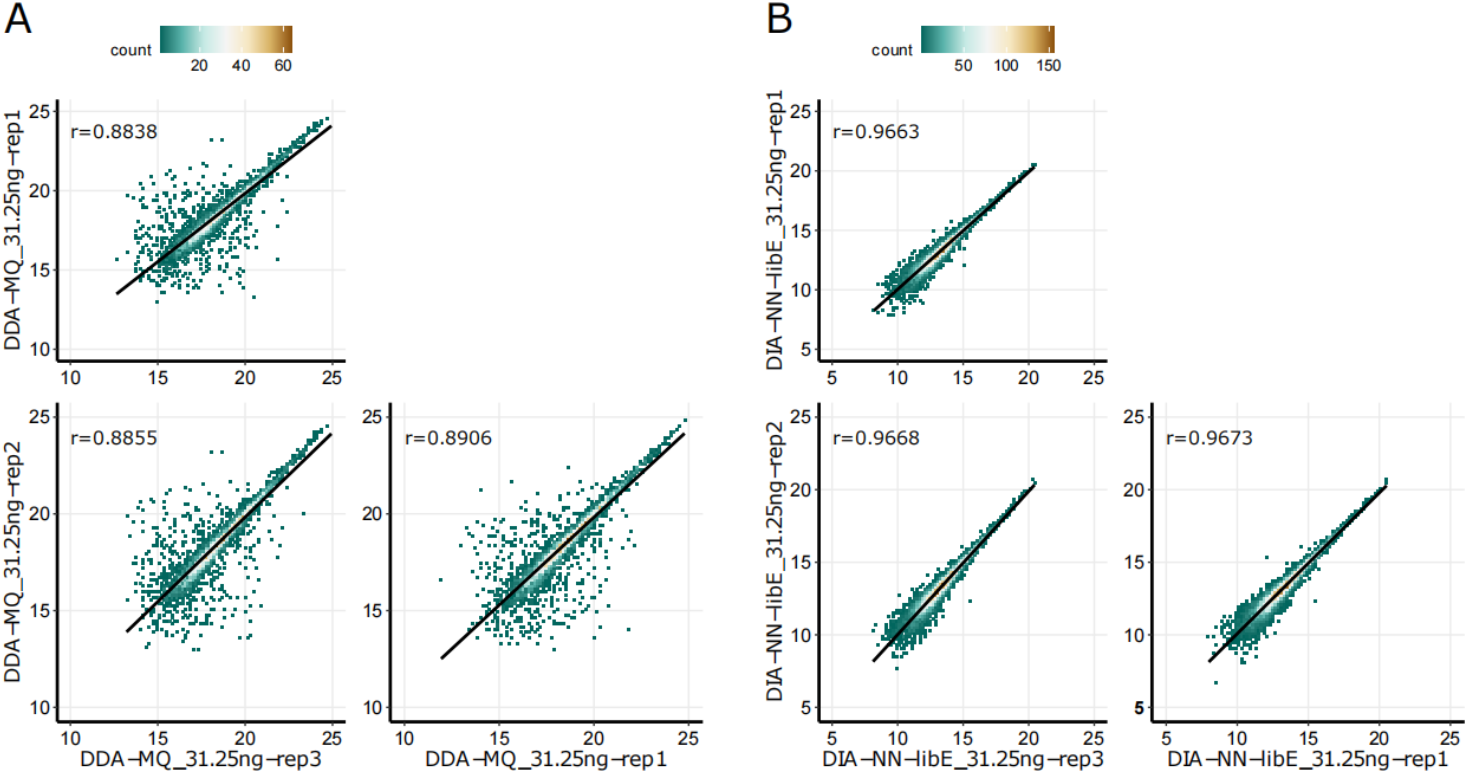
Correlation plots between 3 technical replicates at the lowest peptide quantity tested when using the DDA-MQ (A) and the DIA-NN-libE (B) workflow.

**Supplementary Figure 4:**
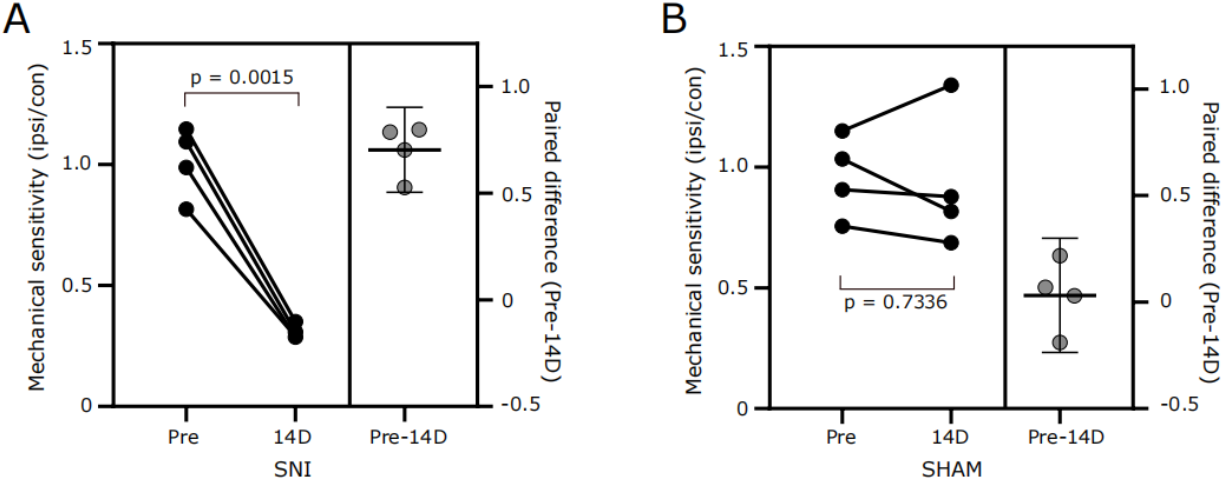
Mechanical sensitivity tests of SNI mice (A) and SHAM mice (B) before (Pre) and after surgery (14D). Each data point on the left side of both graphs corresponds to the ratio of ipsilateral (operated) and contralateral (non-operated) paw values of individual mice (two-tailed paired t-test; N = 4 mice/condition). The right side of both graphs shows the mean (SNI=0.702, SHAM=0.031) and individual differences between Pre and 14D with 95% confidence interval.

**Supplementary Figure 5:**
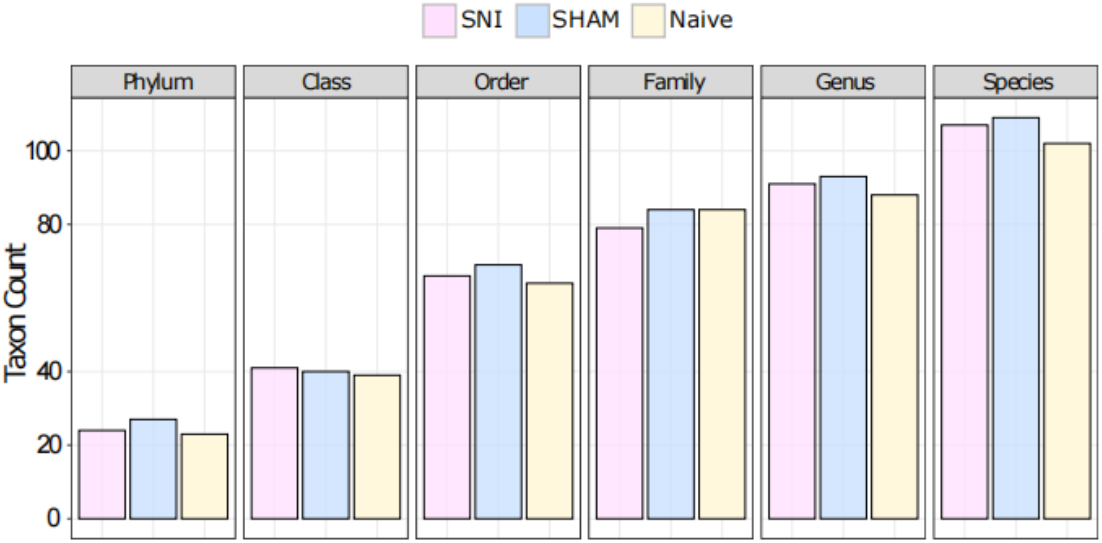
Taxonomic annotations in all three conditions at 6 taxa levels. Cutoff: at least 3 unique peptides per taxon (as identified by iMetalab).

**Supplementary Figure 6:**
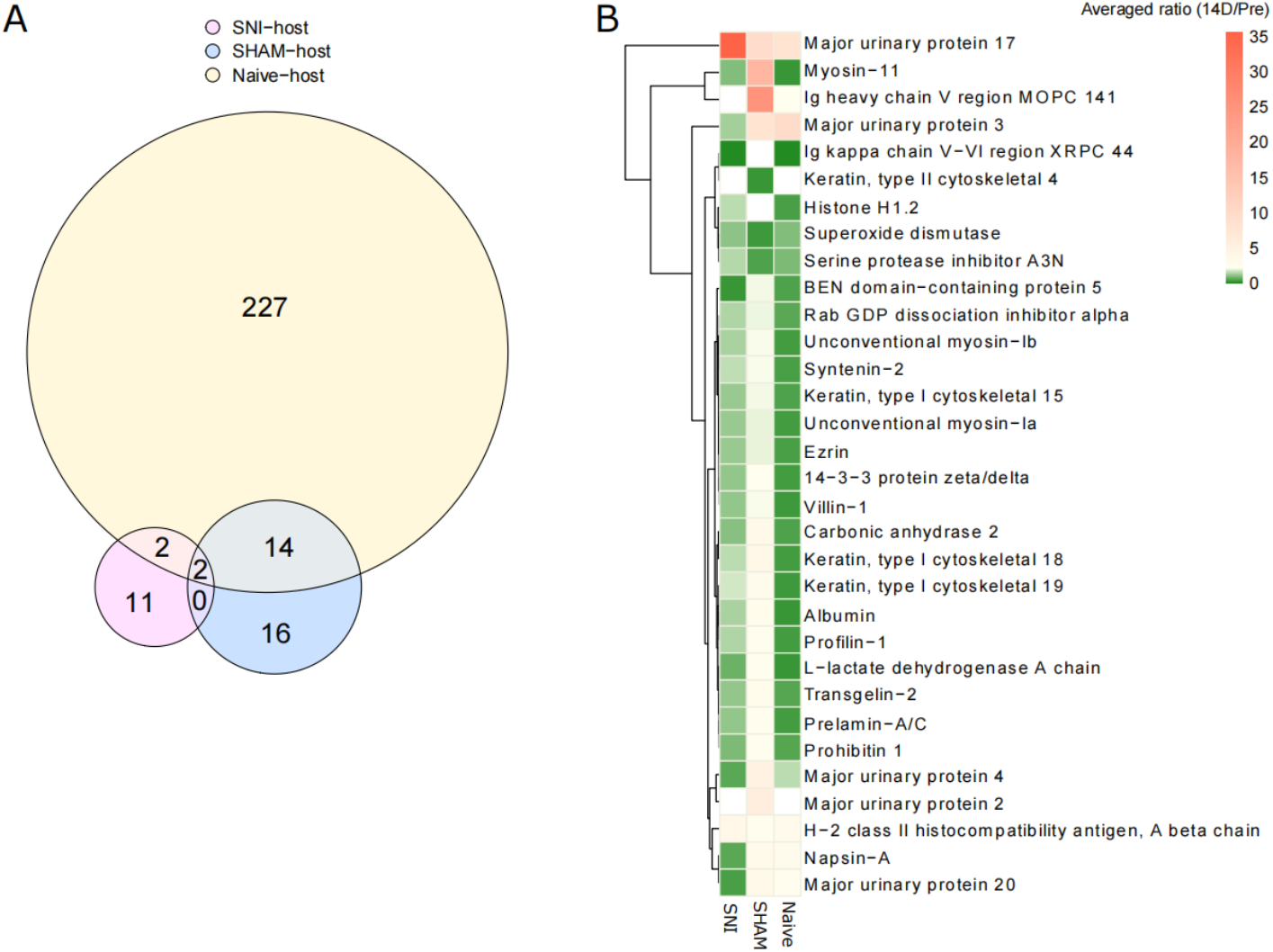
Differential expression in the host proteome. A) Shared and unique numbers of host proteins after pairwise comparison in each condition (p-value < 0.005). B) The expression patterns of 32 host proteins (selected based on p-value < 0.005, pair-wise comparison, 14D versus Pre in SHAM condition) in all three experimental groups. Color code indicates the average ratio (14D/Pre) of each host protein.

## References

1. Prescott, S.L., Wegienka, G., Logan, A.C., and Katz, D.L. (2018). Dysbiotic drift and biopsychosocial medicine: how the microbiome links personal, public and planetary health. Biopsychosoc Med 12, 7. 10.1186/s13030-018-0126-z.

2. Clemente, J.C., Ursell, L.K., Parfrey, L.W., and Knight, R. (2012). The impact of the gut microbiota on human health: an integrative view. Cell 148, 1258–1270. 10.1016/j.cell.2012.01.035.

3. Liu, Y., Beyer, A., and Aebersold, R. (2016). On the Dependency of Cellular Protein Levels on mRNA Abundance. Cell 165, 535–550. 10.1016/j.cell.2016.03.014.

4. Sun, S., Jones, R.B., and Fodor, A.A. (2020). Inference-based accuracy of metagenome prediction tools varies across sample types and functional categories. Microbiome 8, 46. 10.1186/s40168-020-00815-y.

5. Mills, R.H., Vazquez-Baeza, Y., Zhu, Q., Jiang, L., Gaffney, J., Humphrey, G., Smarr, L., Knight, R., and Gonzalez, D.J. (2019). Evaluating Metagenomic Prediction of the Metaproteome in a 4.5-Year Study of a Patient with Crohn’s Disease. mSystems 4. 10.1128/mSystems.00337-18.

6. Fetzer, I., Johst, K., Schawe, R., Banitz, T., Harms, H., and Chatzinotas, A. (2015). The extent of functional redundancy changes as species’ roles shift in different environments. Proc Natl Acad Sci U S A 112, 14888–14893. 10.1073/pnas.1505587112.

7. Tian, L., Wang, X.W., Wu, A.K., Fan, Y., Friedman, J., Dahlin, A., Waldor, M.K., Weinstock, G.M., Weiss, S.T., and Liu, Y.Y. (2020). Deciphering functional redundancy in the human microbiome. Nat Commun 11, 6217. 10.1038/s41467-020-19940-1.

8. Stamboulian, M., Canderan, J., and Ye, Y. (2022). Metaproteomics as a tool for studying the protein landscape of human-gut bacterial species. PLoS Comput Biol 18, e1009397. 10.1371/journal.pcbi.1009397.

9. Duan, H., Cheng, K., Ning, Z., Li, L., Mayne, J., Sun, Z., and Figeys, D. (2022). Assessing the Dark Field of Metaproteome. Anal Chem 94, 15648–15654. 10.1021/acs.analchem.2c02452.

10. Michalski, A., Cox, J., and Mann, M. (2011). More than 100,000 detectable peptide species elute in single shotgun proteomics runs but the majority is inaccessible to data-dependent LC-MS/MS. J Proteome Res 10, 1785–1793. 10.1021/pr101060v.

11. Armengaud, J. (2023). Metaproteomics to understand how microbiota function: The crystal ball predicts a promising future. Environ Microbiol 25, 115–125. 10.1111/1462-2920.16238.

12. Aakko, J., Pietila, S., Suomi, T., Mahmoudian, M., Toivonen, R., Kouvonen, P., Rokka, A., Hanninen, A., and Elo, L.L. (2020). Data-Independent Acquisition Mass Spectrometry in Metaproteomics of Gut Microbiota-Implementation and Computational Analysis. J Proteome Res 19, 432–436. 10.1021/acs.jproteome.9b00606.

13. Long, S., Yang, Y., Shen, C., Wang, Y., Deng, A., Qin, Q., and Qiao, L. (2020). Metaproteomics characterizes human gut microbiome function in colorectal cancer. NPJ Biofilms Microbiomes 6, 14. 10.1038/s41522-020-0123-4.

14. Meier, F., Brunner, A.D., Frank, M., Ha, A., Bludau, I., Voytik, E., Kaspar-Schoenefeld, S., Lubeck, M., Raether, O., Bache, N., et al. (2020). diaPASEF: parallel accumulation-serial fragmentation combined with data-independent acquisition. Nat Methods 17, 1229–1236. 10.1038/s41592-020-00998-0.

15. Xian, F., Sondermann, J.R., Gomez Varela, D., and Schmidt, M. (2022). Deep proteome profiling reveals signatures of age and sex differences in paw skin and sciatic nerve of naive mice. Elife 11. 10.7554/eLife.81431.

16. Meier, F., Beck, S., Grassl, N., Lubeck, M., Park, M.A., Raether, O., and Mann, M. (2015). Parallel Accumulation-Serial Fragmentation (PASEF): Multiplying Sequencing Speed and Sensitivity by Synchronized Scans in a Trapped Ion Mobility Device. J Proteome Res 14, 5378–5387. 10.1021/acs.jproteome.5b00932.

17. Demichev, V., Messner, C.B., Vernardis, S.I., Lilley, K.S., and Ralser, M. (2020). DIA-NN: neural networks and interference correction enable deep proteome coverage in high throughput. Nat Methods 17, 41–44. 10.1038/s41592-019-0638-x.

18. Kolmeder, C.A., Salojarvi, J., Ritari, J., de Been, M., Raes, J., Falony, G., Vieira-Silva, S., Kekkonen, R.A., Corthals, G.L., Palva, A., et al. (2016). Faecal Metaproteomic Analysis Reveals a Personalized and Stable Functional Microbiome and Limited Effects of a Probiotic Intervention in Adults. PLoS One 11, e0153294. 10.1371/journal.pone.0153294.

19. Meier, F., Park, M.A., and Mann, M. (2021). Trapped Ion Mobility Spectrometry and Parallel Accumulation-Serial Fragmentation in Proteomics. Mol Cell Proteomics 20, 100138. 10.1016/j.mcpro.2021.100138.

20. Meier, F., Brunner, A.D., Koch, S., Koch, H., Lubeck, M., Krause, M., Goedecke, N., Decker, J., Kosinski, T., Park, M.A., et al. (2018). Online Parallel Accumulation-Serial Fragmentation (PASEF) with a Novel Trapped Ion Mobility Mass Spectrometer. Mol Cell Proteomics 17, 2534–2545. 10.1074/mcp.TIR118.000900.

21. Cox, J., and Mann, M. (2008). MaxQuant enables high peptide identification rates, individualized p.p.b.-range mass accuracies and proteome-wide protein quantification. Nat Biotechnol 26, 1367–1372. 10.1038/nbt.1511.

22. Demichev, V., Szyrwiel, L., Yu, F., Teo, G.C., Rosenberger, G., Niewienda, A., Ludwig, D., Decker, J., Kaspar-Schoenefeld, S., Lilley, K.S., et al. (2022). dia-PASEF data analysis using FragPipe and DIA-NN for deep proteomics of low sample amounts. Nat Commun 13, 3944. 10.1038/s41467-022-31492-0.

23. Kieser, S., Zdobnov, E.M., and Trajkovski, M. (2022). Comprehensive mouse microbiota genome catalog reveals major difference to its human counterpart. PLoS Comput Biol 18, e1009947. 10.1371/journal.pcbi.1009947.

24. Wang, J., Lang, T., Shen, J., Dai, J., Tian, L., and Wang, X. (2019). Core Gut Bacteria Analysis of Healthy Mice. Front Microbiol 10, 887. 10.3389/fmicb.2019.00887.

25. Treede, R.D., Rief, W., Barke, A., Aziz, Q., Bennett, M.I., Benoliel, R., Cohen, M., Evers, S., Finnerup, N.B., First, M.B., et al. (2015). A classification of chronic pain for ICD-11. Pain 156, 1003–1007. 10.1097/j.pain.0000000000000160.

26. Price, T.J., and Gold, M.S. (2018). From Mechanism to Cure: Renewing the Goal to Eliminate the Disease of Pain. Pain Med 19, 1525–1549. 10.1093/pm/pnx108.

27. Morreale, C., Bresesti, I., Bosi, A., Baj, A., Giaroni, C., Agosti, M., and Salvatore, S. (2022). Microbiota and Pain: Save Your Gut Feeling. Cells 11. 10.3390/cells11060971.

28. Decosterd, I., and Woolf, C.J. (2000). Spared nerve injury: an animal model of persistent peripheral neuropathic pain. Pain 87, 149–158. 10.1016/S0304-3959(00)00276-1.

29. Segelcke, D., Fischer, H.K., Hutte, M., Dennerlein, S., Benseler, F., Brose, N., Pogatzki-Zahn, E.M., and Schmidt, M. (2021). Tmem160 contributes to the establishment of discrete nerve injury-induced pain behaviors in male mice. Cell Rep 37, 110152. 10.1016/j.celrep.2021.110152.

30. Cobos, E.J., Nickerson, C.A., Gao, F., Chandran, V., Bravo-Caparros, I., Gonzalez-Cano, R., Riva, P., Andrews, N.A., Latremoliere, A., Seehus, C.R., et al. (2018). Mechanistic Differences in Neuropathic Pain Modalities Revealed by Correlating Behavior with Global Expression Profiling. Cell Rep 22, 1301–1312. 10.1016/j.celrep.2018.01.006.

31. Chu, M., and Zhang, X. (2022). Bacterial Atlas of Mouse Gut Microbiota. Cellular Microbiology 2022, 5968814. 10.1155/2022/5968814.

32. Beresford-Jones, B.S., Forster, S.C., Stares, M.D., Notley, G., Viciani, E., Browne, H.P., Boehmler, D.J., Soderholm, A.T., Kumar, N., Vervier, K., et al. (2022). The Mouse Gastrointestinal Bacteria Catalogue enables translation between the mouse and human gut microbiotas via functional mapping. Cell Host Microbe 30, 124–138 e128. 10.1016/j.chom.2021.12.003.

33. Szyrwiel, L., Sinn, L., Ralser, M., and Demichev, V. (2022). Slice-PASEF: fragmenting all ions for maximum sensitivity in proteomics. bioRxiv, 2022.2010.2031.514544. 10.1101/2022.10.31.514544.

34. Distler, U., Łącki, M.K., Startek, M.P., Teschner, D., Brehmer, S., Decker, J., Schild, T., Krieger, J., Krohs, F., Raether, O., et al. (2023). midiaPASEF maximizes information content in data-independent acquisition proteomics. bioRxiv, s2023.2001.2030.526204. 10.1101/2023.01.30.526204.

35. Skowronek, P., Krohs, F., Lubeck, M., Wallmann, G., Itang, E.C.M., Koval, P., Wahle, M., Thielert, M., Meier, F., Willems, S., et al. (2023). Synchro-PASEF Allows Precursor-Specific Fragment Ion Extraction and Interference Removal in Data-Independent Acquisition. Mol Cell Proteomics 22, 100489. 10.1016/j.mcpro.2022.100489.

36. Hua, D., Li, S., Li, S., Wang, X., Wang, Y., Xie, Z., Zhao, Y., Zhang, J., and Luo, A. (2021). Gut Microbiome and Plasma Metabolome Signatures in Middle-Aged Mice With Cognitive Dysfunction Induced by Chronic Neuropathic Pain. Front Mol Neurosci 14, 806700. 10.3389/fnmol.2021.806700.

37. Braundmeier-Fleming, A., Russell, N.T., Yang, W., Nas, M.Y., Yaggie, R.E., Berry, M., Bachrach, L., Flury, S.C., Marko, D.S., Bushell, C.B., et al. (2016). Stool-based biomarkers of interstitial cystitis/bladder pain syndrome. Sci Rep 6, 26083. 10.1038/srep26083.

38. Engevik, M.A., Morra, C.N., Roth, D., Engevik, K., Spinler, J.K., Devaraj, S., Crawford, S.E., Estes, M.K., Kalkum, M., and Versalovic, J. (2019). Microbial Metabolic Capacity for Intestinal Folate Production and Modulation of Host Folate Receptors. Front Microbiol 10, 2305. 10.3389/fmicb.2019.02305.

39. Taverner, T., Crowe, F.L., Thomas, G.N., Gokhale, K., Thayakaran, R., Nirantharakumar, K., and Rajabally, Y.A. (2019). Circulating Folate Concentrations and Risk of Peripheral Neuropathy and Mortality: A Retrospective Cohort Study in the U.K. Nutrients 11. 10.3390/nu11102443.

40. Botez, M.I., Peyronnard, J.M., Bachevalier, J., and Charron, L. (1978). Polyneuropathy and folate deficiency. Arch Neurol 35, 581–584. 10.1001/archneur.1978.00500330029005.

41. Mölzer, C., Wilson, H.M., Kuffova, L., and Forrester, J.V. (2021). A Role for Folate in Microbiome-Linked Control of Autoimmunity. Journal of Immunology Research 2021, 9998200. 10.1155/2021/9998200.

42. Bethea, J.R., and Fischer, R. (2021). Role of Peripheral Immune Cells for Development and Recovery of Chronic Pain. Frontiers in Immunology 12. 10.3389/fimmu.2021.641588.

43. Ustianowska, K., Ustianowski, L., Machaj, F., Goracy, A., Rosik, J., Szostak, B., Szostak, J., and Pawlik, A. (2022). The Role of the Human Microbiome in the Pathogenesis of Pain. Int J Mol Sci 23. 10.3390/ijms232113267.

44. Ding, W., You, Z., Chen, Q., Yang, L., Doheny, J., Zhou, X., Li, N., Wang, S., Hu, K., Chen, L., et al. (2021). Gut Microbiota Influences Neuropathic Pain Through Modulating Proinflammatory and Anti-inflammatory T Cells. Anesth Analg 132, 1146–1155. 10.1213/ANE.0000000000005155.

45. Jiang, J., Natarajan, K., and Margulies, D.H. (2019). MHC Molecules, T cell Receptors, Natural Killer Cell Receptors, and Viral Immunoevasins-Key Elements of Adaptive and Innate Immunity. Adv Exp Med Biol 1172, 21–62. 10.1007/978-981-13-9367-9_2.

46. Abokor, A.A., McDaniel, G.H., Golonka, R.M., Campbell, C., Brahmandam, S., Yeoh, B.S., Joe, B., Vijay-Kumar, M., and Saha, P. (2021). Immunoglobulin A, an Active Liaison for Host-Microbiota Homeostasis. Microorganisms 9. 10.3390/microorganisms9102117.

47. Weis, A.M., and Round, J.L. (2021). Microbiota-antibody interactions that regulate gut homeostasis. Cell Host Microbe 29, 334–346. 10.1016/j.chom.2021.02.009.

48. Mirpuri, J., Raetz, M., Sturge, C.R., Wilhelm, C.L., Benson, A., Savani, R.C., Hooper, L.V., and Yarovinsky, F. (2014). Proteobacteria-specific IgA regulates maturation of the intestinal microbiota. Gut Microbes 5, 28–39. 10.4161/gmic.26489.

49. Hughes, C.S., Moggridge, S., Muller, T., Sorensen, P.H., Morin, G.B., and Krijgsveld, J. (2019). Single-pot, solid-phase-enhanced sample preparation for proteomics experiments. Nat Protoc 14, 68–85. 10.1038/s41596-018-0082-x.

50. Xiao, L., Feng, Q., Liang, S., Sonne, S.B., Xia, Z., Qiu, X., Li, X., Long, H., Zhang, J., Zhang, D., et al. (2015). A catalog of the mouse gut metagenome. Nature Biotechnology 33, 1103–1108. 10.1038/nbt.3353.

51. Muth, T., Vaudel, M., Barsnes, H., Martens, L., and Sickmann, A. (2010). XTandem Parser: an open-source library to parse and analyse X!Tandem MS/MS search results. Proteomics 10, 1522–1524. 10.1002/pmic.200900759.

52. Barsnes, H., and Vaudel, M. (2018). SearchGUI: A Highly Adaptable Common Interface for Proteomics Search and de Novo Engines. J Proteome Res 17, 2552–2555. 10.1021/acs.jproteome.8b00175.

53. Vaudel, M., Burkhart, J.M., Zahedi, R.P., Oveland, E., Berven, F.S., Sickmann, A., Martens, L., and Barsnes, H. (2015). PeptideShaker enables reanalysis of MS-derived proteomics data sets. Nature Biotechnology 33, 22–24. 10.1038/nbt.3109.

54. Cuklina, J., Lee, C.H., Williams, E.G., Sajic, T., Collins, B.C., Rodriguez Martinez, M., Sharma, V.S., Wendt, F., Goetze, S., Keele, G.R., et al. (2021). Diagnostics and correction of batch effects in large-scale proteomic studies: a tutorial. Mol Syst Biol 17, e10240. 10.15252/msb.202110240.

55. Cox, J., Hein, M.Y., Luber, C.A., Paron, I., Nagaraj, N., and Mann, M. (2014). Accurate proteome-wide label-free quantification by delayed normalization and maximal peptide ratio extraction, termed MaxLFQ. Mol Cell Proteomics 13, 2513–2526. 10.1074/mcp.M113.031591.

56. Cheng, K., Ning, Z., Zhang, X., Li, L., Liao, B., Mayne, J., and Figeys, D. (2020). MetaLab 2.0 Enables Accurate Post-Translational Modifications Profiling in Metaproteomics. J Am Soc Mass Spectrom 31, 1473–1482. 10.1021/jasms.0c00083.

57. Ritchie, M.E., Phipson, B., Wu, D., Hu, Y., Law, C.W., Shi, W., and Smyth, G.K. (2015). limma powers differential expression analyses for RNA-sequencing and microarray studies. Nucleic Acids Research 43, e47–e47. 10.1093/nar/gkv007.

58. Zhou, Y., Zhou, B., Pache, L., Chang, M., Khodabakhshi, A.H., Tanaseichuk, O., Benner, C., and Chanda, S.K. (2019). Metascape provides a biologist-oriented resource for the analysis of systems-level datasets. Nat Commun 10, 1523. 10.1038/s41467-019-09234-6.

